# Tissue adaptation: implications for gut immunity and tolerance

**DOI:** 10.1101/125641

**Authors:** Ana M. C. Faria, Bernardo S. Reis, Daniel Mucida

## Abstract

Tissue adaptation is an intrinsic component of immune cell development, influencing both resistance to pathogens and tolerance. Chronically stimulated surfaces of the body, in particular the gut mucosa, are the major sites where immune cells traffic and reside. Their adaptation to these environments requires constant discrimination between natural stimulation coming from harmless microbiota and food, and pathogens that need to be cleared. This review will focus on the adaptation of lymphocytes to the gut mucosa, a highly specialized environment that can help us understand the plasticity of leukocytes arriving at various tissue sites and how tissue-related factors operate to shape immune cell fate and function.

## Introduction

“Adaptation (…) could no longer be considered a static condition, a product of a creative past, and became instead a continuing dynamic process.” (Mayr, 1982)

In biology, adaptation generally refers to the process that enhances the fitness of individuals equipped with plasticity, in response to imposed conditions (Mayr, 1982). In this sense, immune cells can be considered highly adaptive entities. First, they display inter- and intratissue migratory capacity. Second, they generally leave primary lymphoid organs in a low-differentiated stage and their final commitment and acquisition of effector functions are determined by interactions with cells and signals in peripheral lymphoid and non-lymphoid organs. Therefore, tissue adaptation is an intrinsic component of immune cell development, influencing both resistance to pathogens and inflammation-induced tissue damage.

In order to perform their critical role in maintaining organismal homeostasis in a continuously changing environment, immune cells circulate extensively even in tissues initially thought to be “immune-privileged” (Shechter et al., 2013). Establishment of tissue-resident immune cell populations enables a quicker response to local stress, injury, or infection. Tissue-resident cells can then further recruit precursors or mature immune cells that participate in the initiation, effector phase and resolution of the inflammatory process, which is highly dependent on the nature of the initial insult, as well as on the target tissue and existing resident immune cells (Medzhitov, 2008).

The surfaces of the body are the major sites where immune cells traffic and reside. The intestinal mucosa alone harbors more lymphocytes than all lymphoid organs combined (Crago et al., 1984; Cerf-Bensussan et al., 1985; van der Heijden, 1986; Guy-Grand et al., 1991a). These tissues pose numerous challenges to recruited immune cells since they are chronically stimulated by a plethora of external agents including microbiota, dietary components, environmental noxious substances and infectious pathogens. Adaptation of immune cells to the intestinal environment requires constant discrimination between the natural stimulation coming from harmless microbiota and food, and pathogens that need to be cleared. Chronic immune activation can lead to tissue injury and proliferation-induced senescence or cancer. Immune cells at the intestinal mucosa therefore must maintain careful control over the balance between inflammation and tolerance. This review will focus on the adaptation of immune cells to the gut mucosa as an example of how tissue environment shapes leukocyte fate and function.

## Tissue-imprinting on mature lymphocytes

Early lymphocyte lineage commitment steps that occur in the primary immune organs (e.g. B versus T cell lineage commitment) are thought to be irreversible under steady-state conditions. Expression of Notch-induced TCF-1 in the thymus, for instance, is a crucial step leading to T cell lineage commitment and Notch-guided T cell receptor (TCR) rearrangement represents an irreversible checkpoint in αβ versus γδ specification since it involves DNA recombination (Weber et al., 2011). Further checkpoints during thymic αβ T cell development are dependent on the interplay between the transcription factors ThPOK/MAZR/GATA-3 and Runx3, leading to mature CD4 and CD8 lineage specification, respectively (Sawada et al., 1994; Siu et al., 1994; Ellmeier et al., 1997; Taniuchi et al., 2002; He et al., 2005; He et al., 2008; Muroi et al., 2008; Setoguchi et al., 2008; Sakaguchi et al., 2010). Similar to αβ- and γ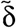specification, αβ T cell CD4- and CD8-MHC (I and II, respectively) restriction is irreversible post-commitment. Although differentiation of mature immune cells into activated effector cells is generally associated with a reduction in their plasticity potential, recent reports indicate that upon migration to tissues these cells undergo further specialization in response to particular environmental conditions **(Figure 1)**.

**Figure 1.**
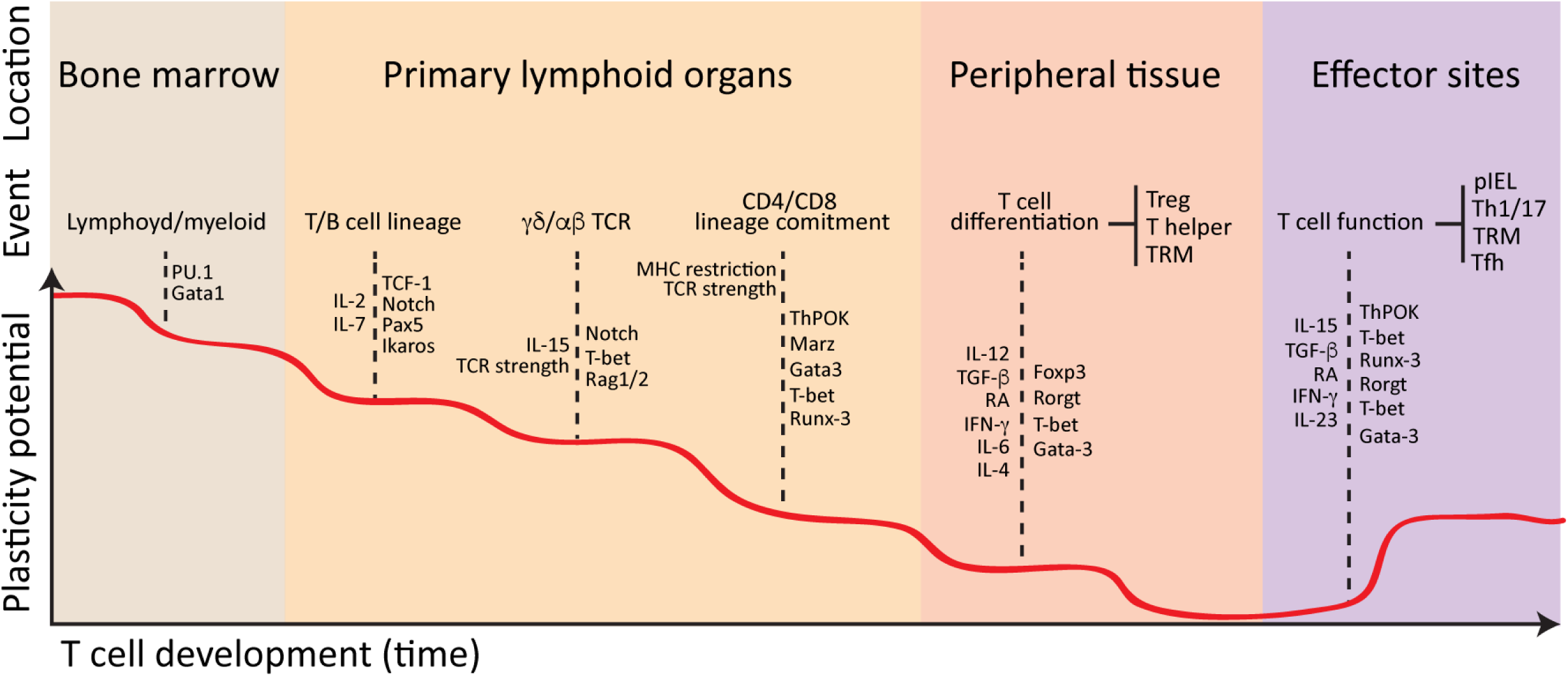
T cell plasticity during lineage commitment. Lymphoid precursors exit the bone marrow and migrate to the thymus where they differentiate into mature T cells. Rag1/2-dependent TCR rearrangement gives rise to TCRγδ and TCRαβ lineages while MHC restriction and TCR strength leads to CD4 or CD8 commitment of the TCRαβ lineage in a Runx3- and ThPOK-dependent manner. Mature CD4 and CD8 T cells exit the thymus and receive further activation and differentiation signals in secondary lymphoid organs and nonlymphoid tissues. Each commitment step is generally associated with loss of cell plasticity, although migration to particular tissue or effector sites may allow re-acquisition of various levels of plasticity, depending on the target tissue, contexts and intra-tissue microenvironments.

During primary immunization or infections, activated effector T cells differentiate into memory T cells with distinct phenotypes, plasticity and functions, depending on which lymphoid or non-lymphoid tissues they seed (Schenkel and Masopust, 2014). Tissue-resident memory (TRM) T cells represent a recently identified T cell population that resides in non-lymphoid tissues without recirculating and with a distinct core gene signature (Mackay et al., 2013; Schenkel and Masopust, 2014). TRM cells are derived from effector T cell precursors and can be recruited to several tissues, particularly barrier surfaces, even in the absence of overt inflammation; albeit inflammatory processes can significantly increase TRM cell differentiation from effector T cell precursors or recruitment to sites such as skin epidermis, vaginal epithelium, lung airways, salivary glands and ganglia (Masopust et al., 2001; Mackay et al., 2013; Shin and Iwasaki, 2013; Laidlaw et al., 2014; Schenkel et al., 2014; Mackay et al., 2016).

Upon migration to the tissue environment, T cells are subject to cues that can promote remarkable plasticity and functional specialization. For instance, epidermal TRM cells adopt a dendritic morphology resembling Langerhans cells, which allows them to probe the epidermal layer and maintain a slow migrating behavior, whereas dermis T cells bearing the same TCR specificity, maintain the typical lymphoid shape, moving considerably faster than epidermal TRM cells (Zaid et al., 2014). Additionally, trafficking of effector T cells is facilitated by secretion of the chemokine CCL25 (ligand for CCR9) in the small intestine, and CCL28 (ligand for CCR10) in the colon as well as the integrin MAdCAM-1 (ligand for α4β7) expressed by intestinal endothelium (Iijima and Iwasaki, 2015). Additional signals secreted by epithelial or resident hematopoietic cells, such as IL-15 and TGF-β, further regulate T cell accumulation and/or retention in the tissue. Rapid, TGF-β-dependent, upregulation of the integrin αEβ7 (CD103, which binds to E-cadherin on epithelial cells) by “memory precursor effector” CD8^+^ T cells was shown to regulate accumulation of TRMs in the intestinal epithelium (Sheridan et al., 2014). Induced expression of certain cell surface molecules, such as CRTAM (class I major histocompatibility complex–restricted T cell–associated molecule) and 2B4, a NK-related receptor, may also be important in the accumulation or maintenance of T cells in the gut epithelial layer (Cortez et al., 2014).

### Adaptation to the intestinal epithelium

Intraepithelial lymphocytes (IELs) primarily reside at the epithelial layer of mucosal surfaces and skin, displaying both innate and adaptive characteristics (Cheroutre et al., 2011). IELs comprise a heterogeneous group of lymphocytes, including cells considered TRM and innate lymphoid cells, and are characterized by high expression levels of activation markers such as CD69; gut-homing integrins; NK-inhibitory and activating receptors such as Ly49 and KIR families; cytotoxic T lymphocyte (CTL)-related genes such as granzyme B; and anti-inflammatory or inhibitory receptors like LAG-3 (Denning et al., 2007a). Another common characteristic of IELs is the surface expression of CD8αα homodimers, which can bind both to classical MHC-I and to epithelial cell-associated non-classical MHC-I molecules (Guy-Grand et al., 1991b; Gangadharan et al., 2006; Cheroutre and Lambolez, 2008; Cheroutre et al., 2011; Guy-Grand et al., 2013).

Similar to Foxp3-expressing Tregs (Fan and Rudensky, 2016)(discussed below), IELs can be divided into peripheral (p) or thymic (t) IELs, depending on their origin and process of differentiation. tIELs are represented mostly by TCRγδ^+^CD8αα^+^ and TCRαβ^+^CD8αα^+^ cells, which migrate shortly after birth from the thymus to the intestinal epithelial layer where their maintenance depends on T-bet- and IL-15-dependent pathways (Guy-Grand et al., 1991a; Lefrancois, 1991; Suzuki et al., 1997; Ma et al., 2009; Malamut et al., 2010; Huang et al., 2011; Mucida et al., 2013; Klose et al., 2014; Reis et al., 2014). However, as evidenced by their migration to the intestinal epithelium prior to microbial colonization, tIELs do not require microbiota for induction or maintenance and their numbers are not altered in germ-free (GF) animals (Bandeira et al., 1990; Mota-Santos et al., 1990). Nevertheless, dietary metabolites, particularly aryl-hydrocarbon receptor (AHR) ligands found in cruciferous vegetables, appear to regulate TCRγδ IEL maintenance in the gut epithelium (Li et al., 2011). Additionally, absence of intact dietary proteins has been shown to impact pIEL numbers (Menezes Jda et al., 2003), suggesting that both tIELs and pIELs continuously require lumen-derived signals for their development or maintenance.

pIELs are comprised of mature CD4^+^ and CD8^+^ TCRαβ^+^ cells t h at m igrate t o th e gut epithelium upon activation in secondary lymphoid tissues and acquisition of gut homing receptors (Das et al., 2003; Masopust et al., 2006; Mucida et al., 2013; Reis et al., 2013; Luda et al., 2016). Contrary to tIELs, which show reduced frequency in ageing animals, pIELs accumulate with age and are severely reduced in GF mice (Mota-Santos et al., 1990; Umesaki et al., 1993; Mucida et al., 2013). In a similar fashion to peripheral Tregs, initial tissue imprinting on pIELs likely takes place in the gut-draining lymph nodes, where TGF-β-and retinoic acid-producing dendritic cells (DCs), primarily IRF8-dependent migratory DCs, induce gut-homing capacity in naïve T cells (Iwata et al., 2004; Coombes et al., 2007; Mucida et al., 2007; Sun et al., 2007; Konkel et al., 2011; Esterhazy et al., 2016; Luda et al., 2016). In contrast to pTregs however, the second step of pIEL tissue imprinting (acquisition of an “IEL phenotype”) takes place in the intestinal tissue, likely within the epithelium itself (Sujino et al., 2016). Like tIELs, T-bet upregulation downstream of IL-15, IFN-γ and IL-27 signaling is required for epithelial imprinting on pIELs (Klose et al., 2014; Reis et al., 2014). Additionally, the intra-tissue modulation of T-box transcription factors T-bet and Eomes was shown to play an essential role in the maturation and maintenance of lung and skin TRM cells, suggesting a broad role for these transcriptional regulators in lymphocyte tissue imprinting (Mackay et al., 2015).

Irrespective of their MHC restriction and TCRαβ lineage specification, both CD4^+^ and CD8^+^ pIELs progressively acquire CD8αα expression and tIEL markers in the gut epithelium, partially reverting their lineage program set during thymic development. This feature is particularly remarkable in CD4^+^ pIELs (CD4-IELs), which induce post-thymic downmodulation of the T helper lineage commitment transcriptional factor ThPOK that is preceded by the induction of both T-bet and Runx3, the latter a CD8 lineage commitment transcription factor (Mucida et al., 2013; Reis et al., 2013; Reis et al., 2014). This epithelium-specific regulation of T cell lineage transcription factors results in a broad suppression of programming associated with mature CD4^+^ T cells, including expression of costimulatory molecules, T helper cytokines and T regulatory-associated transcription factor Foxp3 (Mucida et al., 2013; Reis et al., 2013; Reis et al., 2014; Sujino et al., 2016). The exact mechanisms by which CD4^+^ T cells undergo such drastic functional adaptation towards the IEL program remain to be defined. Nevertheless, in sharp contrast to all other known peripheral CD4^+^ T cells, CD4-IELs acquire CD8αα expression and hence the capacity to engage class I MHC or thymus leukemia antigen (TL), expressed on the surface of gut epithelial cells (Hershberg et al., 1990; Wu et al., 1991; Leishman et al., 2001). Peripheral CD8^+^ T cells also progressively upregulate CD8αα homodimers in the process of tissue adaptation, a feature that allows these cells to further differentiate into long-lived memory cells, and to respond to epithelium-specific challenges (Huang et al., 2011). Regardless of the subset in which it is expressed, CD8αα decreases antigen sensitivity of the TCR-negatively regulating T cell activation (Cheroutre and Lambolez, 2008). Therefore, both CD4^+^ and CD8^+^ pIELs acquire features that distinguish these cells from other peripheral T cells and likely allow them to quickly respond to lumen- or epithelium-specific cues, although a detailed characterization of pIEL populations in additional mucosal sites is yet to be performed. Intriguingly, similar epithelial imprinting has been described in an innate lymphocyte population, which lacks expression of rearranged TCR or CD4/CD8, but does express several of the hallmarks observed in gut epithelial T cells including CD8αα (Van Kaer et al., 2014).

Although IELs exhibit cytotoxic potential, their killing activity is kept in check by tissue imprinting. A positive regulator of the cytotoxic activity of IEL populations is IL-15, a cytokine produced by a wide range of cells including epithelial, stromal and several myeloid cell populations (Jabri and Abadie, 2015). IL-15 is upregulated in several chronic inflammatory diseases, acting both as a local danger signal that promotes Th1 cell-mediated immunity and as a costimulatory signal to effector cytotoxic T cells (Jabri and Abadie, 2015). In contrast to CD8αα expression, which increases the TCR activation threshold of IELs, IL-15 reduces it, hence promoting their lymphokine-activated killer activity (LAK activity) (Meresse et al., 2004; Meresse et al., 2006; Tang et al., 2009; Ettersperger et al., 2016). While physiological levels of IL-15 contribute to the development of intestinal T cells and provide a complementary stress signal to trigger protective CTL functions during intracellular infections, uncontrolled IL-15 production during inflammation can result in T-cell mediated disruption of the epithelial barrier and promote disorders such as coeliac disease (Meresse et al., 2004; Meresse et al., 2006; Tang et al., 2009; DePaolo et al., 2011; Jabri and Abadie,2015). An analogous process was described in the skin epidermis and lung, where intra-tissue modulation of T-bet expression allowed for continuous expression of CD122 (IL-15Rβ) by CD103^+^ TRM cells, promoting effector function and IL-15-dependent survival (Mackay et al., 2013; Mackay et al., 2015). In contrast to both skin and lung TRM cells as well as to CD8αα+ IELs, virus-specific CD8αβ+ TRM cells were reported to be maintained within intestinal mucosa independently of IL-15 (Schenkel et al., 2016). This observation may indicate that IL-15 dependence may vary not only according to the anatomic location, but also to the nature of tissue resident lymphocytes.

In summary, the epithelial milieu tightly adjusts the phenotype of its resident lymphocytes, as evidenced by the strikingly similar gene programs acquired by intestinal IELs, irrespective of their T cell receptor or coreceptor expression, lineage or subtype (Denning et al., 2007a). Although an understanding of the physiological roles played by IELs, or their role in controlling pathogen invasion and intestinal inflammation still requires further studies, several lines of evidence point to their function in maintaining the epithelial cell barrier and responding to pathogens (Boismenu and Havran, 1994; Lepage et al., 1998; Chen et al., 2002; Poussier et al., 2002; Das et al., 2003; Meresse et al., 2006; Olivares-Villagomez et al., 2008; Ismail et al., 2009; Tang et al., 2009; Ismail et al., 2011; Edelblum et al., 2015; Sujino et al., 2016). These studies highlight examples of T cell adaptation to the single-layered intestinal epithelium, a uniquely challenging location for both resistance and tolerance given its close proximity to the highly stimulatory gut lumen.

### Tissue-dependent immune regulation of Tregs

Tregs are also prone to tissue conditioning and represent a noteworthy example of late adaptation to specific environmental signals. Similar to other T cell subsets, pTregs acquire tissue-homing capacity during their differentiation or activation (Fan and Rudensky, 2016). Recent reports indicated that highly suppressive breast tumor-infiltrating Tregs shared a similar gene program to Tregs from normal breast tissue, but not to other peripheral activated Treg cells (De Simone et al., 2016; Plitas et al., 2016). Mechanisms employed by pTregs to prevent or suppress overt inflammation vary depending on the target tissue. For instance, Treg-specific ablation of IL-10 or IL-10 receptor does not result in systemic inflammatory diseases but rather induces severe inflammation in mucosal tissues and in the skin (Rubtsov et al., 2008; Chaudhry et al., 2011; Huber et al., 2011).

The capacity of tissue-resident Tregs to sense particular inflammatory cues is paramount for their regulation of associated responses. This idea is also supported by several studies targeting Treg transcriptional programs associated with specific pathological cues. T-bet expression by Tregs is required for their suppression of T-bet dependent effector T cell responses. Hence, mice carrying T-bet-deficient Tregs develop severe Th1 inflammation including lymphadenopathy and splenomegaly, while other responses are still kept in check (Koch et al., 2009). Additionally, Treg-specific ablation of Stat3, a key transcription factor for Th17 cell differentiation, results in uncontrolled Th17 inflammation in the intestine (Chaudhry et al., 2009). However, ablation of transcription factors IRF4 or GATA-3, which are required for Th2 differentiation in Tregs, results in selective and spontaneous development of allergic- as well as Th17 responses at mucosal tissues and skin (Zheng et al., 2009; Cretney et al., 2011; Wang et al., 2011; Wohlfert et al., 2011). Conversely, conditional deletion of *Rorc*, associated with Th17 development, in Tregs results in increased mucosal Th2 immunity, suggesting that during inflammation Treg sensing of the tissue milieu might reestablish equilibrium by regulating additional arms of the immune system (Ohnmacht et al., 2015; Eberl, 2016). Both Tregs and Treg-inducing signals can also be overturned by certain signals from inflamed tissues. While retinoic acid (RA) is described to induce Treg development (Mucida et al., 2007), in the presence of IL-15, RA was shown to rapidly trigger mucosal DCs to release the proinflammatory cytokines IL-12p70 and IL-23, inhibiting pTreg differentiation while promoting differentiation of Th1 cells and cytotoxic T cell function (DePaolo et al., 2011; Hall et al., 2011). These effector T cell subtypes can then induce tissue-damage in response to luminal antigens, as in gluten intolerance responses observed in coeliac disease patients (DePaolo et al., 2011).

The reported plasticity of the Treg lineage during inflammatory or tissue-specific responses does not result in instability of the Foxp3-dependent Treg program in non-lymphopenic settings, as suggested by fate-mapping strategies using dual reporter strains (Rubtsov et al., 2010). Nevertheless, studies performed under lymphopenic or inflammatory settings raised the possibility that uncommitted Treg populations, preferentially pTregs, lose Foxp3 and in some cases even differentiate into effector T cells (Komatsu et al., 2009; Tsuji et al., 2009; Zhou et al., 2009; Miyao et al., 2012). These studies suggest that some level of instability or plasticity can exist even in committed Treg populations. Using the tamoxifen-inducible *Foxp3*-driven Cre recombinase strain to fate-map bona fide Tregs, we confirmed previous reports that Tregs show stable Foxp3 expression over time in all tissues examined (Rubtsov et al., 2010), including the intestinal lamina propria (Sujino et al., 2016). The only exception to this rule was observed in the intestinal epithelium, where we found that roughly 50% of former Tregs physiologically lose Foxp3 over a period of 5 weeks, in a microbiota-dependent manner (Sujino et al., 2016). These former Tregs developed an IEL phenotype, including acquisition of Runx3 and loss of ThPOK, as described above (Sujino et al., 2016). Perhaps one of the clearest demonstrations of tissue adaptation of Tregs was reported recently in a transnuclear mouse strain generated by somatic cell nuclear transfer from a single pTreg. The authors found that CD4^+^ T cells carrying a monoclonal naturally-occurring pTreg TCR preferentially generated pTregs in gut draining LNs, but almost exclusively produced CD4-IELs in the intestinal epithelium, both in a microbiota-dependent manner (Bilate et al., 2016).

It is currently thought that Tregs utilize several redundant and complementary mechanisms to suppress inflammatory responses and studies described above emphasize the role of environmental sensing in Treg function. The instability of the Treg lineage within the intestinal epithelium may represent an important modulation of regulatory activity that is coordinated by this particular environment (Hooper and Macpherson, 2010; Atarashi et al., 2011; Josefowicz et al., 2012; Bollrath and Powrie, 2013; Furusawa et al., 2013). However, a specific role for CD4-IELs (described above) in cytolytic responses against intracellular pathogens or in triggering inflammation remains to be defined (Mucida et al., 2013; Jabri and Abadie, 2015). An interesting possibility is that these “potentially cytotoxic” CD4^+^ T cells could play a crucial role in the immune responses against viral infections tropic for MHC-II target cells, including HIV-1–infected human CD4^+^ T cells (Khanna et al., 1997) or viruses that managed to escape conventional cytotoxic CD8^+^ T cell-dependent surveillance, including cytomegalovirus and HIV-1 itself (Ko et al., 1979; Simon et al., 2015). Nevertheless, the observation that tissue sensing of particular environmental cues, such as dietary or microbiota metabolites, results in coordinated immune regulation, represents an important step in the understanding of diseases triggered by uncontrolled inflammation.

### Gut imprinting on antibody production

Tissue adaptation of B cells in the gut mucosa also involves cross-talk between these cells and the luminal factors influencing their differentiation. For instance, germ-free mice show reduced levels of secretory IgA while serum IgM levels are preserved relative to SPF animals (Benveniste et al., 1971). Conversely, polyreactive IgA secretion is fundamental to maintenance of microbiota diversity, as evidenced by dysbiosis and additional phenotypic variations observed in IgA-deficient mice (Suzuki et al., 2004; Fransen et al., 2015; Moon et al., 2015). Two distinct populations of B cells can be found in the intestinal lamina propria: “conventional” B2 cells and B1 cells. Both populations undergo differentiation into IgA-producing plasma cells, a hallmark of mucosal sites. B2 cells originate in the bone marrow and undergo isotype switching into IgA^+^ B cells in Peyer’s patches (Craig and Cebra, 1971), cecal patches (Masahata et al., 2014) and isolated lymphoid follicles (ILFs) (Macpherson et al., 2000; Fagarasan et al., 2001; He et al., 2007). A small population of IgA^+^ B1 cells is found exclusively in the lamina propria and the origin of this population is yet to be confirmed, although early reports suggested they arise from peritoneal B1 cells (Beagley et al., 1995; Bao et al., 1998) in a microbiota-dependent fashion (Ha et al., 2006).

Several features of the gut mucosa seem to influence the preferential differentiation of gut B cells towards IgA-producing cells (reviewed elsewhere (Fagarasan et al., 2010; Cerutti et al., 2011; Pabst et al., 2016; Reboldi and Cyster, 2016)). First, in response to microbiota, intestinal follicular T helper cells (Tfh), which provide help for class switch recombination (CSR) and affinity maturation of B cells in the germinal centers of Peyer’s patches (Fagarasan et al., 2002; Kawamoto et al., 2012) preferentially arise from differentiated Th17 and secrete large amounts of TGF-β and IL-21, cytokines associated with IgA class switching (Seo et al., 2009; Hirota et al., 2013; Cao et al., 2015). One example of the importance of this microbiota-induced IgA is the finding that bacterial species known to strongly induce Th17 responses, such as segmented filamentous bacteria (SFB), overgrow in IgA-deficient mice (Suzuki et al., 2004; Fransen et al., 2015). Second, T cell-independent IgG- and IgA-class switching (Macpherson et al., 2000; Bergqvist et al., 2006) ensures innate-like tissue imprinting in B cells, which can be triggered by microbial recognition via TLRs expressed on gut B cells (Koch et al., 2016) or non-hematopoietic cells (Tsuji et al., 2008). In addition to microbial products, sensing of gut factors such as retinoic acid by follicular dendritic cells, which in turn secrete TGF-β and the B cell-activating factor of the TNF family (BAFF), has also been shown to trigger class switch recombination into IgA^+^ B cells (Suzuki et al., 2010). Finally, B1 cells are also associated with T cell-independent IgA class switching in the lamina propria, although this site of B cell commitment to IgA is still debated (Macpherson et al., 2000; Fagarasan et al., 2001). There is a large body of evidence supporting the pre-commitment of plasma cells to IgA in Peyer’s patches, as well as other peripheral sites including spleen, mLNs, ILFs and peritoneal cavity, before their recruitment to the intestinal LP (de Andres et al., 2007; Koch et al., 2009; Lindner et al., 2015; Reboldi et al., 2016). A “post-recruitment adaptation” (i.e. CSR) of B1 cells into IgA-producing plasma cells within the lamina propria has been proposed based on detection of activation-induced cytidine deaminase (AID) expression and DNA excision circles in LP B cells (Macpherson, A. J. et al, 2000; Fagarasan et al, 2001). It still remains possible, however, that this process primarily takes place in ILFs interspersed in the mucosal region (Bergqvist et al., 2006). Another issue is that most of the factors involved in CSR to IgA are also required for IgA plasma cell survival in the LP, which confounds loss-of-function studies. Therefore, further confirmatory studies are required to demonstrate the existence of a post-recruitment adaptation of B1 cells in the gut lamina propria.

Unlike peripheral lymphoid organs where plasma cell differentiation occurs in the vicinity of the follicular areas, IgA-plasma cells are imprinted with mucosal-homing properties and migrate to effector niches in the lamina propria, a process dependent on RA and TGF-β signaling (Pabst et al., 2004; Mora et al., 2006; Seo et al., 2013; Lindner et al., 2015). Additionally and similar to the bone marrow, the lamina propria contains specialized niches for plasma cell survival. Both non-hematopoietic cells, such as intestinal epithelial cells (IECs), and hematopoietic cells, including intestinal dendritic cells and eosinophils, were shown to secrete pro-survival factors such IL-6, CXCL12, BAFF, APRIL and thymic stromal lymphopoietin (TSLP) (Jego et al., 2003; He et al., 2007; Xu et al., 2007; Suzuki et al., 2010; Chu et al., 2011; Tezuka et al., 2011; Chu et al., 2014; Jung et al., 2015). Gut-specific factors including TGF-β and RA are therefore associated with a highly diverse set of lumen-derived metabolites and antigens, which contribute to the specific imprinting of gut-associated B cells, though their influence will also vary depending on intra-tissue microenvironments.

## Tissue micro-environments and niches

Organs and tissues are not homogeneous structures, but rather organized spaces with specific niches and microenvironments where different cell types reside and physiological processes take place. The intestinal mucosa show a high degree of architectural complexity and intra-tissue specialization occurs according to anatomical features. For instance, epithelial and sub-epithelial niches host the majority of immune cells and are in closer proximity to luminal stimulation than submucosal and muscularis regions, which in turn are heavily populated by neuronal processes. The microbiota is mostly located in the colon and ileum, whereas soluble antigens are absorbed in the duodenum and jejunum (Mowat and Agace, 2014). Additionally, different microbial species colonize specific regions along the proximal-distal axis and in regards to their proximity to the epithelial layer, while specific dietary components are absorbed at different regions of the intestines. In turn, the cells that comprise the epithelial layer are themselves highly diverse and regionally specialized. For instance, the follicle-associated epithelium (FAE) of Peyer’s patches and ILFs typically lack goblet and Paneth cells, and enteroendocrine cells (de Lau et al., 2012). Paneth cells and M cells are exclusively found in the small intestine whereas goblet cells predominate in the colonic epithelium. These intra-tissue specialized environments are critical for the developmental and functional adaptation of immune cells in the gut **(Figure 2)**.

**Figure 2.**
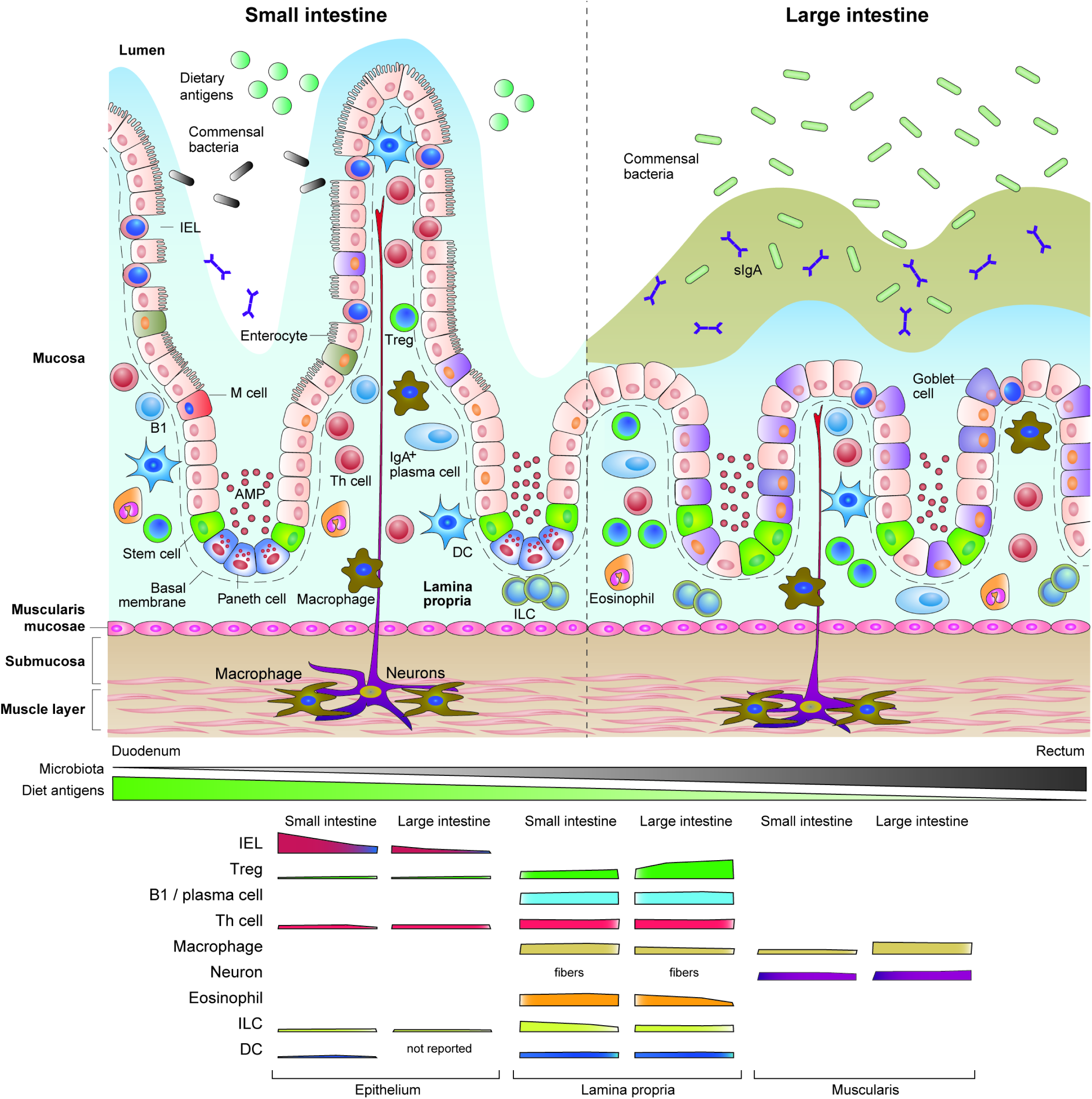
Intestinal microenvironments and niches. The intestine is exposed to constant luminal stimulation and harbors a dense and very diverse set of immune cells. Different layers of the intestinal tissue and regions along the gastro-intestinal tract are subjected to particular stimuli, which are coupled with site-specific adaptation of immune cell subsets. The figure depicts the main characteristics of the intestinal layers, proximal-distal regions and immune cells populations exposed to chronic stimulation by dietary and microbial antigens. While stimulation by dietary antigens or metabolites decreases from proximal to distal intestine, microbial stimulation follows the opposite direction. An approximate illustration of the changes in abundance of each cell type per intestinal region is shown in the lower panel.

### Between the epithelium and lamina propria

Intestinal epithelial cells express sensors for nutrients, microbial antigens and metabolites, as well as for products derived from the interaction between microbes and dietary substances, such as short-chain fatty acids (Hooper and Macpherson, 2010; Shulzhenko et al., 2011; Faria et al., 2013; Derebe et al., 2014). The expression of these various sensing molecules by IECs is tightly regulated by their location along the intestine (e.g. duodenum versus ileum), their positioning within the epithelial layer (e.g. top of the villi versus crypts), their lineage (e.g. Tuft versus enterochromaffin cells), circadian rhythm (Mukherji et al., 2013; Yu et al., 2013) and even by their cellular orientation (basolateral versus apical). These sensing molecules must convey luminal information to neighboring immune cells such as IELs and lamina propria cells as well as to systemic sites (Esterhazy et al., 2016; Loschko et al., 2016). Such IEC-sensing mechanisms are coupled to their role in food absorption, metabolic activities and in their maintenance of surface barriers (Matzinger, 1994; Hooper, 2015). Indeed, recent studies uncovered the impact of microbiota during gestational and pre-weaning periods in the activation and differentiation of IECs and lamina propria cells (Gomez de Aguero et al., 2016; Koch et al., 2016). These effects appeared to be, at least in part, mediated by breast milk- and placenta-derived microbial metabolite-bound antibodies (Gomez de Aguero et al., 2016; Koch et al., 2016), which can be directly sensed by IECs or by underlying lamina propria cells including B cells and ILCs (Gomez de Aguero et al., 2016; Koch et al., 2016; Zeng et al., 2016). These pathways operate at a critical timeframe between birth and weaning by boosting resistance mechanisms against pathogens as well as regulating excessive activation in intestinal immune cells (Verhasselt et al., 2008; Mosconi et al., 2010; Gomez de Aguero et al., 2016; Koch et al., 2016). Therefore, physiological cues derived from the gut lumen at specific life stages support the developmental fitness of the mucosal immune system.

The epithelial layer is separated from the underlying lamina propria by a thin membrane of collagen (basement membrane). Together, the lamina propria and the epithelium harbor the vast majority of immune cells in the body, although the cell populations occupying each layer are strikingly distinct (Mowat and Agace, 2014). In addition, it is also reported that the T cell repertoire differs strongly between the lamina propria and the epithelium (Regnault et al., 1994; Regnault et al., 1996; Arstila et al., 2000; Lathrop et al., 2011; Yang et al., 2014). Consistent with their innate-like phenotype, TCRγδ ⍰⍰⍰ TCRαβ ⍰IELs express restricted oligoclonal T cell repertoires not only in the gut, but also in the skin (Kaufmann, 1996; Probert et al., 2007; Gensollen et al., 2016). Intestinal TCRγδ tIELs express Vγ7 (or Vγ5 depending on the nomenclature utilized) although their ligands are not known and this population seems to be influenced by dietary metabolites found in cruciferous vegetables (AhR ligands), rather than microbiota antigens (Li et al., 2011). The TCR repertoire of double negative (DN) IELs (CD8αα^+^) is also oligoclonal and naturally-occurring DN IEL TCRs are sufficient to induce IEL differentiation when transgenically expressed in developing T cells (Mayans et al., 2014; McDonald et al., 2014). In contrast, the repertoire of colonic lamina propria Tregs is polyclonal and shows little – if any – overlap with the repertoire of naïve or effector CD4^+^ T cells found at the same location, or with other peripheral Tregs (Lathrop et al., 2011). Likewise, the repertoire of intestinal Th17 cells differs significantly from that of other intestinal T cells (Yang et al., 2014). Whereas colonic commensals such as *Clostridium spp*. favor peripheral Treg development (Lathrop et al., 2011; Atarashi et al., 2013), SFB induces differentiation of conventional CD4^+^ T cells into Th17 cells in the distal small intestine (Ivanov et al., 2009; Yang et al., 2014). Additionally, a recent study found a role for dietary antigens in the differentiation of peripheral Tregs occupying the small intestine lamina propria (Kim et al., 2016). Finally, in addition to the divergence of repertoire between lymphocytes occupying the epithelium or lamina propria layers, a study discussed above also indicated that tissue adaptation involves divergence in lineage commitment within the same TCR, since a monoclonal, microbiota antigen-specific TCR elicited strong CD4^+^ pIEL development in the intestinal epithelium, but pTreg induction in the lamina propria (Bilate et al., 2016). Thus, the fate of T cells in the intestine is likely influenced by a combination of their repertoire, luminal stimulation and downstream TCR signaling.

The aforementioned IEL “program” and cell dynamics are unique to the epithelial layer, while the vast majority of macrophages, dendritic cells, ILCs, B cells and Tregs reside in the lamina propria region. Despite differences in their cellular compositions, the epithelial layer constantly influences the state of the lamina propria immune cells. The commensal SFB, which preferentially colonizes the ileum, triggers robust Th17 and IgA responses in the same region (Ivanov et al., 2009). Using complementary strategies to interfere with the bacterium or IECs, studies demonstrated that this adaptive immune response is highly dependent on SFB adhesion to the ileal epithelium (Atarashi et al., 2015; Sano et al., 2015). The authors found that attachment of SFB to the epithelium triggered serum amyloid A (SAA) secretion by IECs. SAA1/2 contributed to both initial Th17 commitment (RORγt expression) in the draining lymph nodes and proliferation of these “poised” Th17 cells (Sano et al., 2015), a process apparently dependent on monocyte-derived CX_3_CR1^+^ cells (Panea et al., 2015).

Reciprocal to the role of IECs in influencing neighboring lamina propria cells, the latter can indeed directly influence the development, proliferation, survival and activity of IECs. ILC subsets, which are segregated into distinct areas of the intestinal mucosa (Sawa et al., 2010; Nussbaum et al., 2013), are particularly relevant players in the crosstalk between the epithelium and lamina propria regions (Cella et al., 2009; Sanos et al., 2009; Buonocore et al., 2010; Geremia et al., 2011; Sawa et al., 2011; Sonnenberg et al., 2012; Fuchs et al., 2013). Besides Th17 differentiation, SFB adhesion was linked to ILC3 production of IL-22 in the lamina propria of the small intestine. ILC3-derived IL-22, in turn, further enhanced IL-17 expression via IEC-derived SAA (Atarashi et al., 2015). Nevertheless, contrary to RORγt^+^ T cells, RORγt^+^ ILCs constitutively secrete IL-22, which is physiologically repressed by IEC-derived IL-25 in a microbiota-dependent manner (Sawa et al., 2011). While IL-22 plays a role in epithelial growth and repair (Lindemans et al., 2015), IFN-γ production by RORγt^+^ ILCs was reported to be required for the protection of the epithelial barrier during enteric infections (Klose et al., 2013).

Several other immune cell types actively participate in the epithelium-lamina propria crosstalk. A subset of B cells has been shown to migrate towards the CCL20 gradient derived from the FAE of the Peyer’s patches, and to influence the differentiation of the specialized M-cells (Golovkina et al., 1999; Ebisawa et al., 2011; Kobayashi et al., 2013). Moreover, in the absence of IgA secretion by lamina propria plasma cells, IECs upregulate resistance mechanisms such as interferon-inducible genes at the expense of their fatty acid processing function, resulting in lipid malabsorption (Shulzhenko et al., 2011). The above studies constitute several parallel examples of coordinated IEC-immune cell responses to luminal perturbations.

### Antigen-presenting cells across intestinal layers

Macrophages and DCs are spread throughout the intestine but display distinct phenotypes and functions depending on the anatomical site they inhabit. A major determinant for such diversity is the ability of migrating monocytes to adapt to specific environmental conditions in the distinct regions of the intestine (Zigmond and Jung, 2013).

Monocyte/macrophage-derived, but not pre-DC-derived, intestinal APC populations express the fractalkine receptor (CX_3_CR1) and these cells constitute the vast majority of intestinal APCs in the mucosal, submucosal and muscularis layers (Gabanyi et al., 2016). Gut macrophage populations play both protective and anti-inflammatory roles, depending on the region they reside or context (Denning et al., 2007b; Bogunovic et al., 2009; Bain et al., 2013; Parkhurst et al., 2013; Bain et al., 2014; Zigmond et al., 2014). Lamina propria macrophages were shown to actively sample luminal bacteria through trans-epithelial dendrites and initiate adaptive immune responses to clear pathogenic bacteria (Rescigno et al., 2001; Niess et al., 2005). On the other hand, these cells were also linked to the initiation and establishment of tolerance to dietary antigens (Hadis et al., 2011; Mazzini et al., 2014).

Pre-DC-derived intestinal (CD103^+^) DCs comprise less than 10% of lamina propria APCs and are virtually absent from the muscularis layer (Bogunovic et al., 2009; Varol et al., 2009; Schreiber et al., 2013). In contrast to other DC populations found in peripheral lymphoid tissues, but similar to other DC populations found in the lung and in the gut-draining lymph nodes, lamina propria DCs constitutively produce RA and TGF-β, which induce gut-homing, differentiation of pTregs and pIELs, and contribute to IgA class switching as discussed above (Mora et al., 2003; Iwata et al., 2004; Johansson-Lindbom et al., 2005; Mora et al., 2006; Coombes et al., 2007; Mucida et al., 2007; Sun et al., 2007; Luda et al., 2016). Similar to gut macrophages, lamina propria DC function has also been associated with triggering effector T cell differentiation and inflammation (Johansson-Lindbom et al., 2005; Laffont et al., 2010; Siddiqui et al., 2010; Semmrich et al., 2012; Persson et al., 2013). However, whereas lamina propria macrophages contact the epithelial layer through their cytoplasmic extensions, CD103^+^ DCs are able to migrate across the basement membrane to the epithelial barrier and sample entire bacteria, a process further enhanced during enteric infections (Farache et al., 2013). Further, while lamina propria macrophages most efficiently sample soluble luminal proteins (Schulz et al., 2009; Farache et al., 2013), epithelial CD103^+^ DCs, but not lamina propria DCs, are also capable of this process (Farache et al., 2013). Therefore, the functional differentiation of dendritic cells and macrophages in the gut mucosa also appears to be a consequence of a regional adaptation process modulated by environmental cues.

Gut macrophages also exhibit a high degree of functional specialization dependent on their proximity to the gut lumen. Lamina propria macrophages preferentially express a pro-inflammatory phenotype whereas macrophages located at the muscularis region, distant from luminal stimulation but nearby neuronal plexuses, display a tissue-protective phenotype that includes genes associated with alternatively activated macrophages (Gabanyi et al., 2016). Moreover, muscularis macrophages were shown to regulate physiological processes such as the basal firing of enteric neurons and peristalsis via secretion of BMP-2 in a microbiota-dependent manner (Muller et al., 2014). Conversely, live imaging indicated that these macrophages are able to sense neuronal signals (Gabanyi et al., 2016). Indeed, commensal colonization was associated with secretion of M-CSF by enteric neurons (Balmer et al., 2014) while enteric infections were linked to the quick activation of extrinsic sympathetic ganglia, which in turn modulate adrenergic β2R^+^ muscularis macrophages to further boost their tissue-protective program (Gabanyi et al., 2016). This intra-tissue adaptation of gut macrophages may have important implications for infection-or inflammation-induced tissue damage (Medzhitov et al., 2012).

## Conclusion

Adaptation, as we propose, is a lifelong process rather than a once-for-all event. Since mucosal surfaces are constantly challenged by fluctuating environmental perturbations, these tissues themselves need to adapt continuously, and recent work has demonstrated that mature immune cells at these sites display a remarkable adaptive capacity. Although progress has been made in defining how immune cells use their plasticity to deal with complex environments such as skin and mucosal surfaces, several questions remain in the field: What are the environmental cues that trigger immune cell adaptation and how do they influence each particular step of lineage commitment? How do changes associated with cellular aging in different tissues influence immune cell adaptation? What are the molecular, including genetic and epigenetic, mechanisms that operate during convergence of distinct immune cell populations within a tissue, or divergence of identical populations in different tissues or under distinct insults? To tackle these questions appropriately, further cross talk between immunology and additional areas of biology, such as neuroscience, metabolism and ageing, is *conditio sine qua non*.

## Acknowledgments

We thank Laura Mackay (The University of Melbourne), Pia Dosenovic (The Rockefeller University), Ainsley Lockhart and additional Mucida lab members for critical reading of this manuscript. A.M.C.F. is supported by a sabbatical internship grant from FAPEMIG, Brazil (ETC-00154-16). B.S.R and D.M. are supported by the Crohn’s & Colitis Foundation of America. D.M. is also supported by the Burroughs Wellcome Fund (PATH award) and by the National Institutes of Health NIH R01 DK093674 and R21 AI131188 grants.

